# Comparison of the effects of running and badminton on executive function: a within-subjects design

**DOI:** 10.1101/625046

**Authors:** Shinji Takahashi, Philip M. Grove

**Author notes:** Corresponding Author: Shinji Takahashi, Address: 2-1-1 Tenjinzawa, Izumi-ku, Sendai, Miyagi, 981-3193 Japan.

## Abstract

Multiple cross-sectional studies have shown that regular complex exercises, which require cognitive demands (e.g., decision making) and various motions, are associated with greater positive effects on executive functions compared to simple exercises. However, the evidence of a single bout of complex exercises is mixed, and investigations on the acute effect of complex exercise using a well-controlled within-subjects research design are few. Therefore, we compared the acute effects of complex exercise on inhibitory functions with those of simple running. Twenty young adults performed three interventions, which were running, badminton, and seated rest as a control condition for 10 min each. During each intervention, oxygen consumption and heart rate were monitored. A Stroop test and a reverse-Stroop test were completed before and after each intervention. The intensities of the badminton and running were equivalent. Badminton significantly improved performance on the Stroop task compared to seated rest; however, running did not enhance performance on the Stroop task relative to seated rest. A single bout of complex exercise elicits a larger benefit to inhibitory function than a single bout of simple exercise. However, the benefit of complex exercise may vary depending on the type of cognitive control.

## Introduction

Regular exercise can prevent cognitive decline and dementia [1]. Moreover, it is thought that exercise has a beneficial effect on executive function, including inhibitory control, working memory, and cognitive flexibility [2]. To clarify what kinds of exercises improve executive functions, researchers have studied both quantitative characteristics (e.g., intensity, duration, and frequency) and qualitative characteristics (e.g., exercise mode and complexity) [3, 4]. Several studies [5–7] showed that complex exercises, including open skill sports (e.g., basketball, tennis, and fencing), have more positive effects on executive functions than simple exercises, such as closed skill sports (e.g., running and swimming). Voss et al. reported results from a meta-analysis indicating that athletes who are experts at complex exercises tend to exhibit superior executive function than simple sports athletes and non-athletes. Complex exercises require the coordination of a variety of motions and cognitive processes, including information pick-up, decision making, visual attention, and inhibition of inappropriate actions.

Given that regular exercise is an aggregation of daily single bout exercises, acute complex exercises might have a different influence on executive functions or activate different brain regions than acute simple exercises. We hypothesized that these features of complex exercises might have effects on executive functions that differ from simple exercises. Specifically, we expected that acute complex exercise would result in a greater benefit to executive function than acute simple exercise.

There is abundant evidence that acute simple aerobic exercise has a significant effect on executive functions [8, 9]. However, the acute effects of complex exercises have received much less attention [10]. Several studies have compared the effects of the different exercise modes on executive functions, however, results from these studies are inconsistent. Studies that support the idea that cognitive demands of complex exercises yield a positive effect on executive functions include Budde et al.[11] who reported that acute coordinative exercise improved selective attention compared to acute simple circuit training. Additionally, Pesce et al. [12] showed that exercise involving a team game improved immediate memory recall function more than a control condition while circuit training failed to show a similar effect. Lastly, Ishihara et al. [13] reported that both playing tennis matches and participating in tennis drills enhanced executive functions relative to a control condition. However, improvement of executive functions following tennis matches was greater than for the drills.

In contrast, other studies have reported that complex exercise impacts executive function to a lesser extent than simple exercise. For example, Gallotta et al. [14] reported that the acute effect of brief basketball mini games on selective attention was smaller than both a running program and a control condition (sitting in an academic class). O’Leary et al. [15] measured inhibitory function after participants walked on a treadmill, played a video game, played an active video game, or sat and rested. The authors reported that walking on a treadmill enhanced inhibitory function compared with playing a videogame and seated rest. Playing active videogame that requires cognitive demands resulted in inhibitory function intermediate to walking and the video game. Kamijo and Abe [16] reported that cycling enhanced executive function while cycling with the cognitive task did not improve executive function but increased cognitive fatigue.

The conflicting evidence outlined above might be due to the methods employed. Many studies use heart rate (HR) as a measure of exercise intensity to equate exercise conditions [11–16]. However, it is known that HR is sensitive to many factors such as gender, exercise mode, emotion, posture, and environmental conditions [17, 18] and so different values for HR may be due to factors other than exercise intensity. To reduce the possibility of the influence of these factors, we included oxygen consumption (VO_2_) and carbon dioxide output (VCO_2_) measures in addition to HR to monitor the intensity of physical activity in our experiment.

Another issue is several previous studies were conducted in a field setting, such as a physical education program or sports training. Experiments in a field setting can be affected by extraneous variables such as weather condition, motivation, day of the week, and anxiety of participants in an unusual situation. To our knowledge, O’Leary et al. [15] and Kamijo and Abe [16] are the only studies on this topic that were conducted in a laboratory setting. Although the exercise tasks in Kamijo and Abe’s experiment were well controlled in terms of intensity, their complex exercise condition was artificial, involving cycling while performing an unrelated cognitive task. In order to resolve the discrepancies in the literature, well-controlled and naturalistic laboratory studies are required. This is the goal of our paper.

To further investigate the acute effects of complex exercises on executive functions, we compared the impact of a single bout of complex exercise on inhibitory function with that of a simple aerobic exercise using a within-subject design employing natural exercises, running and badminton. We monitored exercise intensity via HR, VO_2_, and VCO_2_. We chose Badminton as the complex exercise for this study because it involves various motions such as jumping and racket swinging as well as cognitive demands such as strategy, and shot choice/placement.

We hypothesized that the change of inhibitory function after badminton will be greater than the changes after running. We measured responses to a modified Stroop task and a reverse Stroop task before and after sessions of badminton, running, and seated rest.

## Materials and Methods

### Participants

Sample size was calculated using power analysis for a one-way repeated measures ANOVA with partial eta squared (η_p_^2^) of 0.10, power (1 - β) of 0.80, intraclass correlation coefficient of 0.5 and alpha at 0.05. This analysis indicated that a sample size of 16 was adequate. Participants consisted of undergraduate students from Tohoku Gakuin University who volunteered to participate in the study. A total of 20 healthy participants (8 men, 12 women) were included in the final analysis. All participants were determined to be free of any cardiopulmonary and metabolic disease and visual disorder. The participants were asked to refrain from alcohol use and strenuous physical activity for 24 h before each experiment, and from smoking, food or caffeine consumption for 2 h preceding the experiments. Written informed consent was obtained from all participants before the first experiment. The Human Subjects Committee of Tohoku Gakuin University approved the study protocol. Table 1 shows the characteristics of the participants.

**Table 1.** Characteristics of the participants (mean ± SE).

| Variable | Total ($N = 20$) | Men ($N = 8$) | Women ($N = 12$) |
| --- | --- | --- | --- |
| Age (years) | 20.9 ± 0.2 | 20.6 ± 0.4 | 21.1 ± 0.2 |
| Height (cm) | 164.3 ± 0.4 | 174.9 ± 1.7 | 157.3 ± 1.8 |
| Weight (kg) | 59.8 ± 0.7 | 73.7 ± 0.9 | 50.6 ± 2.0 |
| BMI ($\text{kg} \cdot \text{m}^{-2}$) | 21.9 ± 0.9 | 24.1 ± 0.9 | 20.4 ± 0.6 |
| VO<sub>2</sub>peak ($\text{mL} \cdot \text{kg}^{-1} \cdot \text{min}^{-1}$) | 44.6 ± 1.3 | 50.5 ± 1.7 | 40.7 ± 0.9 |
| HRpeak (bpm) | 197.0 ± 1.5 | 195.8 ± 3.7 | 197.8 ± 1.5 |

### Procedures

#### Day 1

Participants were required to visit the sports physiology laboratory in the gymnasium on four different days (average interval, 5.8 ± 1.4 days). During the first visit, each participant received a brief introduction to this study and completed informed consent. Their height and weight were measured using a stadiometer and a digital scale, respectively. Next, the complete group version of the Stroop/reverse-Stroop color-word test by Hakoda and Sasaki [19] was administered to familiarize participants with the test of inhibitory function. A graded exercise test was subsequently conducted to determine peak of VO_2_ (VO_2_peak) and HRpeak.

#### Day 2-4 (experimental sessions)

Laboratory visits 2, 3, and 4 were experimental sessions. To minimize any order or learning effects, the orders of the experimental sessions were counterbalanced. After arrival at the laboratory, the participants wore a HR monitor (Model RS800cx; Polar Electro Oy, Kempele, Finland) and they rested on a comfortable chair for 10 min. In the experimental sessions, the participants completed the Stroop/reverse-Stroop test (duration: 6 min) before and after each intervention. After the pre-test of Stroop/reverse-Stroop test, the participants were fitted with a portable indirect calorimetry system (MetaMax-3B; Cortex, Leipzig, Germany). This took 1 min and participants rested on a chair for an additional 3 min. For the badminton intervention, the participants moved from the laboratory to a badminton court, which took 2 min. For both the running and the control interventions, the participants walked on a treadmill at 4.2 km·h^-1^ for 2 min, which served as a counterpart to the move from the laboratory to the badminton court. Subsequently, the participants performed each intervention. Based on the protocol of Budde et. al. [11], the duration of the intervention was set to 10 min. After each intervention, the participants returned to the laboratory or walked on the treadmill for 2 min, and then rested for 3 min on a chair. After that, they removed the indirect calorimetry system, completing the post-test of Stroop/reverse-Stroop test, which took 6 min.

In the badminton intervention, the participants played a singles game against one of the two investigators who had experience playing badminton. The investigators played at a level of proficiency that matched the participant’s level and also provided the participants with advice for improvement during the games. During the game, the scores were not recorded and “victory or defeat” was not determined. In the running intervention, the participants ran on a treadmill. Running speed was set according to each participant’s 75%VO_2_peak, which has been previously shown to be the intensity equal to that of the badminton intervention [20]. In the control intervention, the participants were seated on a comfortable chair with their smart phones and were instructed to spend time operating their smartphones as normal. Oxygen consumption, VCO_2_, and HR were monitored throughout each experimental session. Physiological measures for the last 7 min were averaged, and the rating of perceived exertion (RPE) was evaluated at the end of each intervention.

### Aerobic fitness assessment

Participants performed the graded exercise test on a motor-driven treadmill (O2road, Takei Sci. Instruments Co., Niigata, Japan) to volitional exhaustion. The initial speed was set at 7.2 to 9.6 km·h^-1^, according to the estimated physical fitness level of each participant. Each speed lasted 2 min and the speed was increased by 1.2 km·h^-1^ until volitional exhaustion. The portable indirect calorimetry system (MetaMax-3B) measured VO_2_ and VCO_2_, and the average of the final 30 s was defined as the peak oxygen consumption (VO_2_peak). The Polar HR monitor (Model RS800cx) was used to measure HR during the test, and RPE was obtained at the end of each stage. Volitional exhaustion was reached based on the following criterion: 1) RPE ≥ 17, 2) HR ≥ 95% of age-predicted HRmax (220 minus age), and 3) a respiratory exchange ratio (RER VCO_2_· VO_2_^-1^) ≥ 1.10.

### Inhibitory function tasks

We assessed each participant’s inhibitory function by the Stroop/reverse-Stroop test which is composed of a Stroop task and a reverse-Stroop task. The Stroop/reverse-Stroop test is a pencil and paper exercise that requires manual matching rather than oral naming of items. It consists of four subtests arranged in the following order: Test 1 is a neutral task that serves as the control for the reverse-Stroop test. Here, a color name (e.g., red) in black ink is in the leftmost column and five different color patches (red, blue, yellow, green, and black) are placed in right side columns. Participants are asked to check the patch corresponding to the color name. Test 2 is the reverse-Stroop test. Here, a color name (e.g., red) written using a colored ink (e.g., blue) is in the leftmost column and five different color patches are in the right-side columns. Participants are instructed to check the patch corresponding to the color name in the leftmost column. Test 3 is a neutral task that serves as the control for the Stroop test. Here, a color patch (e.g., red) is in the leftmost column and five different color names in black ink are in the right-side columns. Participants are asked to check the color name corresponding to the color patch in the leftmost column. Test 4 is the Stroop test in which a color name (e.g., red) written using a colored ink (e.g., blue) is in the leftmost column and five color names in black ink are in the-right side columns. Participants are instructed to check a word corresponding to the color of the word in the leftmost column. Each subtest consists of 100 items and the materials are printed on an A3-size paper.

The Stroop/reverse-Stroop test includes practice trails (10 items in 10 s) that precede each subtest. In each subtest, participants were instructed to check as many correct items as possible in 60 s. Assessment of inhibitory function was defined as the difference in correct responses between neutral and incongruent tasks. In accordance with Etnier and Chang [21], the performances in Tests 1 and 3 were used as indices of information processing speed and those in Tests 2 and 4 as indices of inhibitory function.

### Statistical analysis

All measurements were described as mean ± standard error. Statistical analyses were conducted using IBM SPSS 25 (SPSS Inc., Chicago, IL, USA). To examine the exercise intensity of each intervention, %HRmax, %VO_2_peak, RER, and RPE were compared using one-way repeated ANOVAs with within-subject factor of mode (running, badminton, and control) and Bonferroni multiple comparison tests separately.

The Stroop tasks (Tests 3 and 4) and reverse-Stroop tasks (Tests 1 and 2) were compared using three-way repeated ANOVAs with within-subject factors of condition (neutral and incongruent), time (pre- and post-test), and mode (running, badminton, and seated rest). When any significant interactions were noted, two-way repeated ANOVAs with within-subject factors of time and mode as post hoc analysis were conducted within each subtest. A significant interaction in two-way repeated ANOVA indicates different changes in performance (pre-test minus post-test) among the interventions. If an interaction was significant, differences in performance changes for each intervention were compared using paired *t* tests. To control for significance level through a series of analyses for the Stroop and reverse-Stroop tasks, the significance levels in each analysis were adjusted by Bonferroni inequality: significance levels of three-way repeated ANOVAs, two-way repeated ANOVAs, and paired *t* tests were set at *p* = .05, *p* = .025, and *p* = .008, respectively. Partial eta squared (η_p_^2^) was calculated as effect size of interactions and main effects in repeated ANOVAs. Cohen’s *d* was also calculated using Bonferroni multiple comparison and paired *t* tests.

## Results

### Intensity of interventions

Table 2 presents the intensities for each intervention. One-way repeated ANOVAs for %VO_2_peak, %HRpeak, RER, and RPE revealed the significant main effects (*F* (2, 38 ≥ 26.4, *p* < .001, η_p_^2^ ≥ 0.58). The % VO_2_peak, %HRpeak, RER, and RPE during both the badminton and running interventions were significantly higher than those during the control intervention (*p* < .001, Cohen’s *d* ≥ 1.40). Differences in all intensity measures between the badminton and running interventions were not significant (*p* ≥ .318, Cohen’s *d* ≤ |0.38|).

**Table 2.** Intensities of each intervention (mean ± SE).

| Variable | Intervention | Total (N = 20) |
| --- | --- | --- |
| %VO<sub>2</sub>peak (%) | Badminton | 76.3 ± 2.1 * |
| | Running | 72.7 ± 1.6 * |
| | Seated rest | 9.5 ± 0.4 |
| %HRpeak (%) | Badminton | 80.9 ± 1.4 * |
| | Running | 81.1 ± 1.8 * |
| | Seated rest | 37.5 ± 1.2 |
| RER (VCO<sub>2</sub> · VO<sub>2</sub><sup>-1</sup>) | Badminton | 0.98 ± 0.01 * |
| | Running | 0.97 ± 0.02 * |
| | Seated rest | 0.85 ± 0.01 |
| RPE | Badminton | 12.9 ± 0.4 * |
| | Running | 13.6 ± 0.5 * |
| | Seated rest | 6.1 ± 0.1 |

\* Significantly different from seated rest; $p < .05$ at Bonferroni multiple comparison tests.

### Stroop task

Table 3 shows the cognitive performances for each intervention. For the Stroop tasks (Tests 3 and 4), three-way repeated ANOVA found a significant interaction between condition, time, and mode (*F* (2, 38) = 4.2, *p* = .022, η_p_^2^ = 0.18). To analyze the significant interaction, two-way repeated ANOVA was conducted separately for Tests 3 and 4. For Test 3, no significant interaction (*F* (2, 38) = 0.9, *p* = .419, η_p_^2^ =. 04) and no significant main effect of mode (*F* (2, 38) = 1.7, *p* = .201, η_p_^2^ = 0.08) were observed; however, a significant main effect of time (*F* (1, 19) = 63.1 *p* < .001, ηp^2^ = .77) was found. For Test 4, a two-way repeated ANOVA revealed a significant interaction (*F* (2, 38) = 5.6, *p* = .007, η_p_^2^ = .23) and a significant main effect of time (*F* (1, 19) = 31.7, *p* < .001, η_p_^2^ =.63); however, a main effect of mode was not significant (*F* (2, 38) = 0.4, *p* = .648, η_p_^2^ =.02). Figure 1 shows the changes in performance for each intervention. As the interaction was significant, differences in the changes in Test 4 for each intervention were compared using the paired t tests. The change in the badminton intervention was significantly greater than that in the control intervention (*t* (19) = 3.6, *p* = .002, Cohen’s *d* = 0.80), while the change in the running intervention was not greater than that in the control (*t* (19) = 1.3, *p* = .207, Cohen’s *d* = 0.29). No difference between the badminton and running interventions was observed (*t* (19) = 1.8, *p* = .082, Cohen’s *d* = 0.44).

**Table 3.** Cognitive performances in each intervention (mean ± SE).

| Task | Condition | Intervention | Pre-test ($N = 20$) | Post-test ($N = 20$) |
| --- | --- | --- | --- | --- |
| Stroop task | Neutral<br>(Test 3) | Badminton | $53.6 \pm 1.2$ | $57.1 \pm 1.4$ |
| | | Running | $55.0 \pm 1.3$ | $57.2 \pm 1.2$ |
| | | Control | $52.6 \pm 1.5$ | $56.0 \pm 1.6$ |
| | Incongruent<br>(Test 4) | Badminton | $48.8 \pm 1.5$ | $53.8 \pm 1.7$ |
| | | Running | $50.3 \pm 1.6$ | $52.9 \pm 1.4$ |
| | | Control | $49.9 \pm 1.8$ | $51.2 \pm 1.7$ |
| Reverse-Stroop task | Neutral<br>(Test 1) | Badminton | $73.0 \pm 1.6$ | $76.9 \pm 1.6$ |
| | | Running | $74.2 \pm 1.3$ | $75.9 \pm 1.8$ |
| | | Control | $71.1 \pm 2.3$ | $74.3 \pm 1.8$ |
| | Incongruent<br>(Test 2) | Badminton | $60.6 \pm 1.7$ | $60.7 \pm 2.3$ |
| | | Running | $61.0 \pm 1.9$ | $60.7 \pm 1.6$ |
| | | Control | $60.3 \pm 2.1$ | $60.5 \pm 1.9$ |

**Fig 1.**
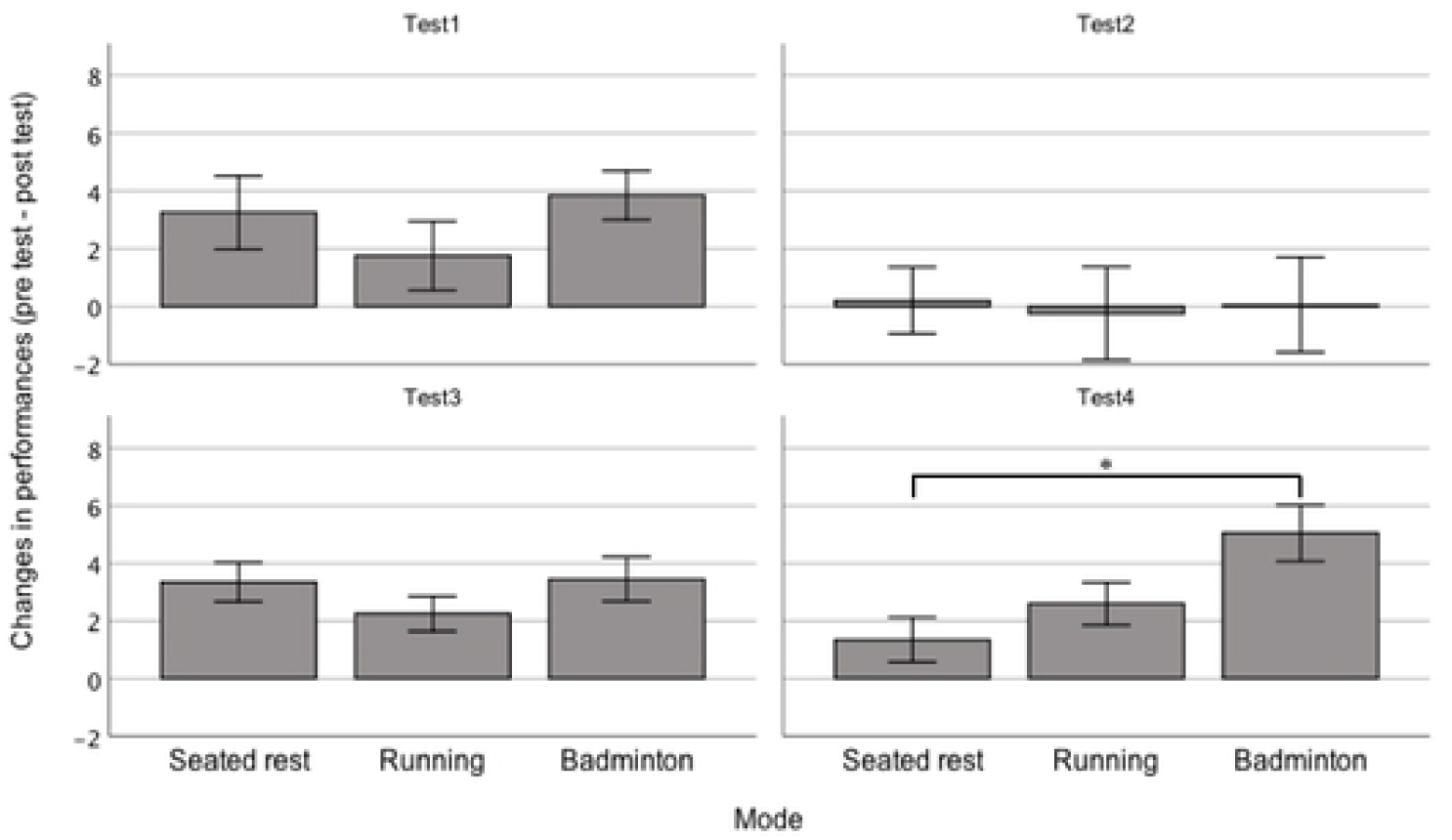
Comparisons of the changes in performances (pre-test minus post-test) between modes in each subtest of the Stroop/reverse-Stroop test. Test 1 is reverse-Stroop neutral test, Test 2 is a reverse-Stroop incongruent test, Test 3 is a Stroop neutral test, and Test 4 is a Stroop incongruent test. Error bars represent standard error. The asterisk (*) indicates a significant difference identified by paired t tests (*p* = .008 adjusted by Bonferroni inequality).

### Reverse-Stroop task

For the reverse-Stroop tasks (Tests 1 and 2), three-way repeated ANOVA found a significant interaction between condition and time (*F* (2, 38) = 8.6, p = .009, η_p_^2^ =.31). To analyze this significant interaction, two-way repeated ANOVAs were conducted for Tests 1 and 2. For Test 1, no significant interaction (*F* (2, 38) = 1.0, *p* = .378, η_p_^2^ =.050) and no significant main effect of mode (*F* (2, 38) = 1.9, *p* = .168, η_p_^2^ = 0.09) were noted; however, a significant main effect of time (*F* (1, 19) = 18.2, *p* < .001, η_p_^2^ = .49) was found. For Test 2, no significant interaction (*F* (2, 38) < 0.1, *p* = .975, η_p_^2^ <. 01) and no significant main effects of mode (*F* (2, 38) = 0.7, *p* = .937, η_p_^2^ < .01) and time (*F* (1, 19) < 0.1, *p* = .999, η_p_^2^ < .01) were observed.

## Discussion

This study aimed to investigate the effect of brief acute complex exercise on inhibitory functions by comparing the effect of badminton with the effect of running on inhibitory function. The main findings of this study were that badminton increased performance in inhibitory function, as shown in the improved performance on the Stroop incongruent test (Test 4), compared to seated rest, while treadmill running did not have a similar effect. Furthermore, changes in performance in the neutral tests (Tests 1 and 3), which served as indices of information processing speed, were not influenced by exercise. These findings indicate that a single bout of complex exercise may selectively improve inhibitory functions compared to simple exercise. However, neither badminton nor running influenced the reverse-Stroop incongruent test (Test 2).

It has been suggested that the reverse-Stroop effect is attributable to brain structures that differ from those in Stroop effects [22, 23]. If this is the case, perhaps, complex exercise impacts brain structures associated with the Stroop effect to a greater extent than those structures associated with the reverse-Stroop effect. Though we cannot make a claim regarding the reverse-Stroop interference, the balance of our results supports our hypothesis that acute complex exercise has a greater effect on executive functions than acute simple exercises.

There were no differences in intensity between the badminton and running interventions, indicating that both interventions were equally categorized as high intensity [24]. In particular, it should be noted that there was no difference of RER (VCO_2_·VO_2_^-1^) between the badminton and running interventions. Statistically equivalent RERs in badminton (0.98 ± 0.01) and running (0.97 ± 0.02) showed that both exercises were the same not only in terms of aerobic energy expenditure but also in anaerobic energy expenditure. Therefore, differences in the effects on cognitive performance between the badminton and running interventions can be attributed to differences in cognitive demand and motions.

The effects of running on changes in performance did not significantly differ from those of control condition in each subtest. Our finding that simple aerobic exercise for 10 min did not benefit cognitive functions is consistent with Chang et al. [8]. These authors reported significant effects of moderate to very high intensity exercise on cognitive functions when the duration of exercise was greater than 11 min. However, the effects of brief exercise of less than 10 min are small and negative. Given Chang et al.’s results, the effect of high intensity running in our study was possibly counteracted by the short exercise duration. Thus, the absence of a significant effect of running on cognitive functions is not unexpected. Furthermore, the effect size (Cohen’s *d* = 0.29) of the running intervention in our study is comparable to those in recently reported meta-analyses [8, 9].

In contrast with running, badminton increased performance compared to seated rest in the Stroop incongruent test (Test 4). Although the pre-intervention versus post-intervention change in the badminton condition did not differ significantly from that of the running condition, the effect size between badminton and running was not small (Cohen’s *d* = 0.44). Changes in performance associated with the running intervention were intermediate between seated rest and badminton (see Fig 1). These results suggest that the cognitive aspects of badminton provide benefits to inhibitory cognitive function over and above the effect of the running. In badminton, players are required to not only grasp the speed and orbit of the shuttle, spatial position of the opponent, but also to choose appropriate shots (e.g., clear, smash, or drop) and perform them. Such cognitive demands could activate the regions of the brain concerned with executive functions. We conclude that the large effect of the badminton intervention on executive function was due to the cognitive demands required to play the game.

Our observation that badminton enhanced inhibitory function to a greater extent than running supported our hypothesis, indicating that the influence of cognitive demands during brief complex exercises is greater than the effects of inefficient exercise. This is consistent with previous studies [11, 12, 25]. However, our observed effects of complex exercise might be restricted to short durations. The effect of complex exercise might be small or negative if exercise duration is extended. Kamijo and Abe [16] reported that 20 min of cycling enhanced executive function while 20 min of cycling with the cognitive task did not improve executive function but increased cognitive fatigue. One possible interpretation of these results is that cognitive fatigue induced by cognitive demand during exercise may cancel the positive effects of acute exercise on executive functions. This interpretation of Kamijo and Abe [16] might explain the results of Gallotta et al [14]. For instance, cognitive demands during exercises might activate the regions of the brain concerned with executive functions (e.g., the prefrontal cortex, anterior cingulate cortex) for a short duration. However, that activation might be gradually overloaded and attenuate the performance of executive functions if exercise duration is extended. This speculation is based on the assumption that complex exercises activate parts of the brain concerned with executive functions more than simple exercises. In order to confirm these assumptions, neuroimaging (e.g., fMRI and fNIRS) and/or electrophysiological evaluations (e.g., ERP P3) are required.

In contrast to the Stroop tasks, we did not observe any differences between badminton and running in the reverse-Stroop tasks. Performance in the reverse-Stroop incongruent test (Test 2) was not influenced by mode, time, or interaction (*p* ≥ .702, η_p_^2^ ≤ .02). This is inconsistent with a few previous studies [26, 27] that have demonstrated that the reverse-Stroop effect is a sensitive index of inhibitory functions for a single bout of exercise. Other study [22] reported that the reverse-Stroop effect differs from the Stroop effect depending on the order the conditions are presented. Furthermore, it has been reported that the Stroop and reverse Stroop effects are mediated by different brain regions [23, 28]. However, the reverse Stroop effect has not been extensively investigated and interpretation of the data is not clear. The mechanisms and the validity of the reverse-Stroop task need further investigations.

One limitation of this study was that the badminton intervention differed from a real badminton match. Victory or defeat was not determined, and the investigators as opponents provided participants tips to improve their game. Therefore, the participants may not have experienced any psychological pressure. In a real badminton match, psychological pressure and stress may influence inhibitory function. Second, none of the participants in this study were experienced badminton players. If well-trained badminton players participated, the observed results may differ. This is because the specific motions and cognitive demands in badminton are overlearned by experienced player and are no longer complex.

## Conclusions

In conclusion, a single bout of a short duration complex exercise selectively enhances inhibitory function relative to a short duration simple exercise. Cognitive demands required in a complex exercise may result in a greater positive effect on executive functions than the negative effect of less efficient motions. However, short duration complex exercise did not improve the performance in the reverse-Stroop incongruent test. The influence of a short duration complex exercise may vary with the type of cognitive tasks.

## Acknowledgements

The author is grateful to all participants and the two badminton instructors. The author also thanks Dr. Keita Kamijo for providing valuable comments.

